# GABAergic signaling promotes early-life seizures in epileptic SYNGAP1^+/-^ mice

**DOI:** 10.1101/2020.11.14.381749

**Authors:** Brennan J. Sullivan, Pavel A. Kipnis, Simon G. Ammanuel, Shilpa D. Kadam

## Abstract

**Objective:** *SYNGAP1* encephalopathy is a developmental and epileptic encephalopathy caused by pathogenic loss of function variants. *Syngap1*-heterozygous (Het^+/-^) mice demonstrate progressive epilepsy with multiple seizure phenotypes in adulthood. Here, we investigate early-life seizures in Het^+/-^ pups and explore of *Syngap1* encephalopathy during development.

**Methods:** Post-natal day 7 (P7) and P12 mice were investigated by tethered video-electroencephalographic (vEEG). The effects of GABAergic drugs phenobarbital (PB) and pentylenetetrazol (PTZ) were investigated at P7 and P12, respectively. 24h tethered vEEG was performed at P24, and telemetric 24h vEEG with 6h sleep deprivation was performed at P35. The effect of perampanel (PMP), an AMPA receptor antagonist, was investigated at P24.

**Results:** Het^+/-^ mice have spontaneous early-life seizures that lack an overt behavioral phenotype. These subclinical seizures are refractory to PB, but the GABA_A_ receptor (GABA_A_R) antagonist PTZ significantly reduced seizure frequency suggesting that GABAergic signaling may promote seizure generation in Het^+/-^ pups. At juvenile ages, Het^+/-^ pups recapitulated the early emergence of high gamma (35-50Hz) during NREM and disruption of behavioral-state gamma homeostasis. This biomarker was significantly exacerbated in Het^+/-^ pups after increasing sleep pressure with sleep deprivation.

**Significance:** Seizures during development have adverse effects on cognitive function. Therefore, an improved understanding of the *SYNGAP1* epilepsy during developmental ages is necessary to delineate the deleterious interactions between aberrant synaptic function and recurrent seizures. The development of evidence-based therapies for early-life intervention will benefit from these insights.

## Introduction

SynGAP is a GTPase-activating protein that plays a major role in the development, structure, and function of excitatory synapses^1^. Pathogenic loss-of-function (LoF) variants in *SYNGAP1* are a leading cause of non-syndromic intellectual disability (ID), autism spectrum disorder (ASD), and epilepsy^2–4^. *SYNGAP1* encephalopathy is a generalized developmental and epileptic encephalopathy (DEE) that includes epilepsy, intellectual disability, severe behavioral problems, ASD, sleep difficulties, and delayed development of speech and motor skills (OMIM# 612621)^5^. Truncating, splice-site, missense mutations, and microdeletions have been reported in patients^5,6^. The majority of pathogenic variants arise from truncating or missense mutations in *SYNGAP1* exons 4-15, which are within the coding region for the core domain of the protein^1,5,6^.

Divergent trajectories of brain maturation have been identified in DEEs, ASD, and other neurodevelopmental disorders^7^. During human cortical development, the expression trajectory of *SYNGAP1* undergoes a robust increase that peaks between birth and the first year of life^8^. In the postnatal cortex, *SYNGAP1* is expressed predominantly in excitatory and inhibitory neurons^9^. SynGAP undergoes splicing at its C-terminus to produce four isoforms that are differentially expressed during development and have distinct effects on synaptic plasticity and dendritic structure^10,11^. The genotype-phenotype correlation for pathogenic *SYNGAP1* variants is currently unresolved, however patients with mutations in exons 1-4 have been associated with a milder ID phenotype^5^. The effects of seizure frequency, age of seizure onset, antiepileptic drug therapy, and interictal EEG abnormalities on the SYNGAP1 phenotypic spectrum are currently unknown.

Heterozygous SynGAP mice (Het^+/-^) present with impaired learning and memory, sensory processing, hyperactivity, sociability, and epilepsy in adulthood^12–19^. During development, Het^+/-^ pups demonstrate precocious unsilencing of thalamocortical synapses, abnormal dendritic spine dynamics, accelerated neuronal maturation, and reduced experience-dependent plasticity^13,20,21^. Previous studies suggest that *Syngap1* has a strong genetic control over synaptic maturation during mouse development^13,20,21^, however the presence of early-life seizures and their subsequent impact is unknown in Het^+/-^ mice.

Previously, we have characterized the natural progression of the epilepsy and seizure phenotypes at advancing adult ages in Het^+/-^ mice^15^. This was associated with a significant impairment in cortical gamma (35-50Hz) and a significant increase in parvalbumin (PV) interneuron GluA2 expression. Since *SYNGAP1*-related DEE presents early in life and the expression of SYNGAP1 is highest during the perinatal period, we investigated the epilepsy in neonatal and juvenile Het^+/-^ mice. The juvenile 24h EEGs combined with sleep deprivation protocols investigated presence of the novel EEG biomarker at the younger age and quantified the effect of increased sleep pressure on the novel biomarker.

## Methods

### Neonatal EEG-Phenobarbital vs. PTZ

As previously described^22^, EEG recordings were acquired using Sirenia Acquisition (version 1.6.4,Pinnacle Technology, Inc.) with synchronized video recording. Data were acquired with 400 Hz sampling rate that had a preamplifier gain of 100, at 0.5–50 Hz. At P7 or P12, pups were implanted with 3 subdermal EEG electrodes (SWE-L25, Ives EEG Solutions): 1 recording and 1 reference overlaying the left/right parietal cortex, and 1 ground overlaying the rostrum while under isoflurane anesthesia (3%-1.5%). Electrodes were fixed in place using cyanoacrylate adhesive. Pups were tethered to a preamplifier inside the recording chamber and allowed to recover from anesthesia (~10min) before continuous video EEG recording in a chamber maintained at 37°C with isothermal pads. For PB experiments, P7 pups were administered a loading dose of PB (25mg/kg, I.P.) after 1h of recording. As previously described^23^, P12 pups were administered PTZ (20mg/kg, I.P) after 1h of recording. At the end of all recordings, the electrodes were removed, and pups were returned to the dam.

### Seizures, IIS, and Behavior Scoring

All scoring was performed by a scorer blinded to genotype and sex. Spontaneous seizures were identified by manual review of all EEGs, as previously described^24^. All seizures were scored within 10s epochs and were defined as ictal events that consisted of rhythmic spikes of high amplitude with a diffuse peak frequency ≥7-8Hz. All IIS were scored within 5s epochs and were defined as high amplitude spikes that were not associated with seizures. Neonatal behavior was scored on video alone with a scorer blind to EEG, genotype, and sex. The behavioral grade of flexor spasms/jerks, waddling, or a behavioral seizure was administered for each 10s epoch.

### 24h vEEG/EMG and PMP Dosing at P24

All surgical procedures and perampanel (PMP) dosing implemented in this study were as previously published^15^. At P18 subdural EEG and suprascapular EMG electrode implantation was performed under isoflurane anesthesia (4%–1.5%). Briefly, subdural EEG and suprascapular EMG electrode implantation was performed under isoflurane anesthesia (4%–1.5%). We used coordinates from bregma for consistent placement of the EEG screw electrodes. After recovering from electrode implantation surgery, mice were placed in a recording chamber with food and water provided ad libitum. For perampanel (PMP) experiments, mice were given a 1mg/kg dose of PMP at 6pm before recording then a second 2mg/kg dose of PMP at 10am.

### Sleep Deprivation at P40

Mice were taken from their home cage and briefly anaesthetized using isoflurane for wireless headcap connection (Pinnacle Technology, 8274-SL). Telemetric EEG recording was enabled, and mice were placed in a 20cm diameter polycarbonate cage with bedding and 18cm steel sleep deprivation rod. A 30min baseline recording was generated after mice awoke from anesthesia. After baseline recordings were made, the Sleep Deprivation Unit (Pinnacle Technology) was activated at speed 3 and mice were sleep deprived for 6h. After 6h, the Sleep Deprivation Unit was inactivated and a 2cm^2^ piece of cloth bedding was placed into the polycarbonate cage alongside food pellets. In the 18h recording period after sleep deprivation, mice had *ad libitum* access to food and water. Telemetric recordings were ended 24h after initiation of sleep deprivation, after which mice were returned to their home cages.

### Western Blotting

All animals for immunochemical characterizations were anesthetized with chloral hydrate (90 mg/ml; IP) before being transcardially perfused with ice-cold saline. The whole fresh brains were removed, the cerebellum was discarded, and the left and right hemispheres were separated. Brains were further micro-dissected into cortex and hippocampus with deep gray matter and stored at −80°C until further processing. Brain tissue homogenates were made and suspended in TPER cell lysis buffer containing 10% protease/phosphatase inhibitor cocktail. Total protein amounts were measured using the Bradford protein assay (Bio-Rad, Hercules, CA, USA) at 570nm and the samples diluted for 50μg of protein in each sample. 20μL of protein samples were run on 4-20% gradient tris-glycine gels (Invitrogen, Gand Island, NY, USA) for 120min at 130V and were transferred onto nitrocellulose membranes overnight at 20V. After the transfer, the nitrocellulose membranes underwent a 1h blocking step in Rockland buffer before 6h incubation with primary antibodies (for all antibody RRIDS, see Key Resources Table): mouse α-KCC2 (1:1000, Millipore), rabbit α-phospho-KCC2-S940 (1:1000 Aviva Systems Biology), rabbit α-phospho-KCC2-T1007 (1:1000; Phospho solutions), rabbit α-NKCC1 (1:1000 Sigma-Aldrich), and mouse α-actin (1:10000, LI-COR Biosciences). Nitrocellulose membranes were then incubated with fluorescent secondary antibodies (1:5000, goat α-rabbit and goat α-mouse, Li-Cor Biosciences, USA). Chemiluminescent protein bands were analyzed using the Odyssey infrared imaging system 2.1 (LI-COR Biosciences). The optical density of each protein sample was normalized to their corresponding actin bands run on each lane for internal control. Mean normalized protein expression levels were then calculated for respective hemispheres.

## Results

### Electrographic seizures in SYNGAP1^+/-^ pups

At P7, continuous 2h vEEG recordings identified the presence of recurrent spontaneous seizures in Het^+/-^ pups (Figure 1A-B). A diverse seizure burden was present in Het^+/-^ pups with paroxysmal EEG activity that included seizures and interictal spiking (IIS; Figure 1B-C). Comparing the frequency of seizures to the frequency of IIS revealed a positive correlation (Figure 1D). Recurrent spontaneous seizures persisted in Het^+/-^ pups to P12 (Figure 1 E-F), no significant differences in seizure frequency were identified between males and females (Figure 1F). To evaluate if seizures were associated with any abnormal behavior in Het^+/-^ pups, video was scored independent of the EEG (Figure 2A). At P7, Het^+/-^ pups had a significantly greater number of flexor spasms/jerks than WT^+/+^ (Figure 2B and Supplemental Video 1). The proportion of seizures that were associated with any concomitant behavior was below 50% (Figure 2C). These seizures were only distinguished by their abnormal EEG patterns and were not associated with the graded (1-3) behaviors during seizures (Figure 2D and Supplemental Video 1). In summary, most early-life seizures were electrographic and required vEEG for identification.

**Figure 1.**
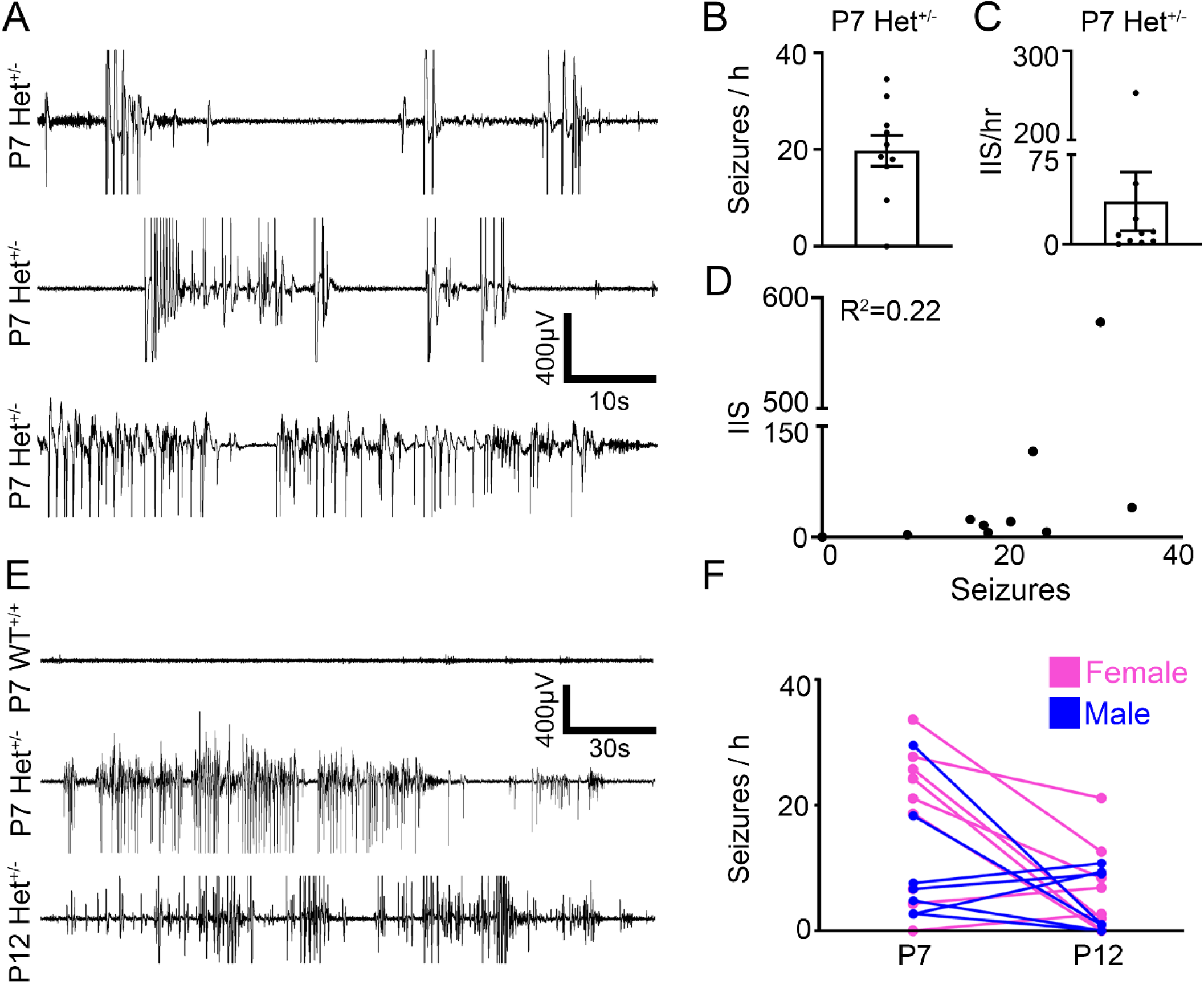
Early Life Seizures in SYNGAP1^+/-^ Pups. (**A**) Representative EEG traces of spontaneous epileptiform discharges in P7 male and female Het mice show bursts of spike wave discharges of variable durations. (**B**) Seizure burden and (**C**) Interictal spikes (IIS) per h at P7 (n=10). (**D**) Correlation of seizure burden (i.e., ictal events >6 sec duration) and IIS at P7. (**E**) EEG traces from WT P7, Het P7, and Het P12 mice. (**F**) Seizures per h at P7 and P12 for Het mice (n=5 male and n=10 female). For 1h P7 WT and P7 Het EEG traces see Supplemental Figure 1.

**Figure 2.**
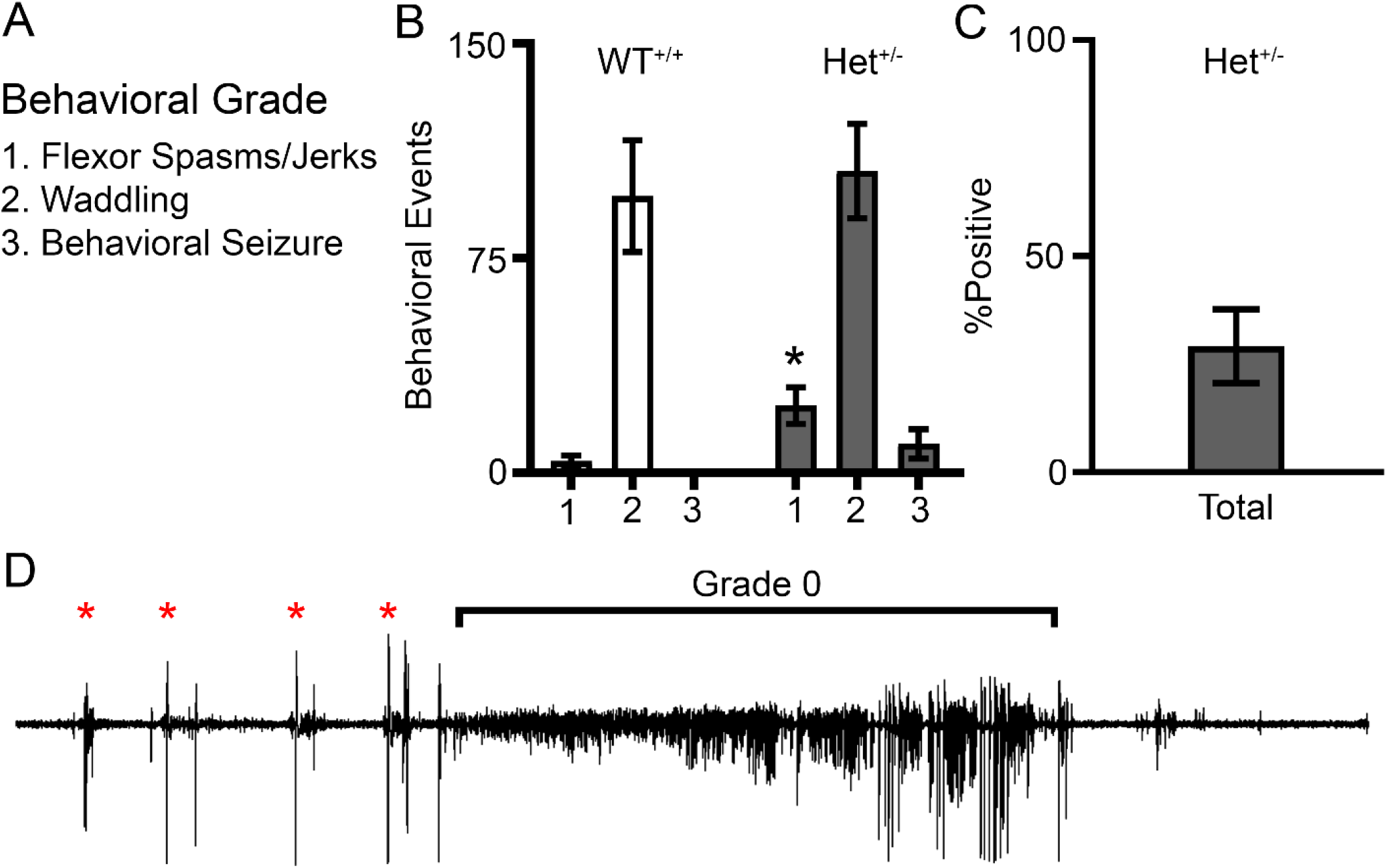
P7 SYNGAP1^+/-^ epileptiform discharges are subclinical. (**A**) Graded behavioral grading parameters on video for P7 neonatal mice during vEEG recording. (**B**) WT and Het behaviors during recording (n=6 WT and n=8 Het). (**C**) Proportion of Het behaviors scored on video only those that were associated with a concomitant epileptiform discharge on EEG. (**D**) Representative Grade 0 epileptiform discharge that was graded as an electrographic only seizure. Representative trace recorded from a Het pup. Red asterisks denote Grade 1 behaviors of flexor spasms/jerks followed by a ~2 min long grade 0 seizure. See Supplemental Video 1 of the same ictal event with synchronous video. *P<0.05 by two-tailed t-test.

### GABAergic signaling is associated with ictogenesis in Syngap1^+/-^ pups

P7 Het pups with high seizure burdens are refractory to a loading dose of PB, a positive allosteric modulator of GABA_A_Rs (Figure 3A). The efficacy of PB as an anti-seizure medication (ASM) is strongly influenced by neuronal Cl^−^ regulation^25^. The ubiquitously expressed Cl^−^ importer NKCC1 and chief neuronal Cl^−^ exporter KCC2 regulate Cl^−^ levels in neurons^25^. Hippocampal and cortical expression of these Cl^−^ cotransporters was investigated to evaluate if differences in their expression could be implicated in the inefficacy of PB to rescue seizures (Figure 3A-B). There was no significant difference between genotypes in KCC2 expression or in the phosphorylation of sites S940^26^ and T1007^27^ (Figure 3D-E). KCC2 expression was significantly greater in the hippocampus of WT^+/+^ pups compared to cortex (P = 0.0008), but not Het^+/-^ pups (P = .2178). Furthermore, the expression of NKCC1 and the ratio of NKCC1/KCC2 were also not significantly different between genotypes (Figure 3F-G). Previously, the GABA_A_R antagonist PTZ was administered to a KCC2 hypofunction mutant mouse model^26^ that transitioned to status epilepticus after PTZ administration^23^. Therefore, PTZ (20mg/kg IP) was administered to P7 Het^+/-^ pups to investigate if a reduction in GABAergic tone could induce status epilepticus (Figure 3H). However, PTZ reduced the frequency of seizure events and the total duration of seizures (Figure 3I-J). These results suggest that GABAergic signaling is ictogenic in Het^+/-^ pups.

**Figure 3.**
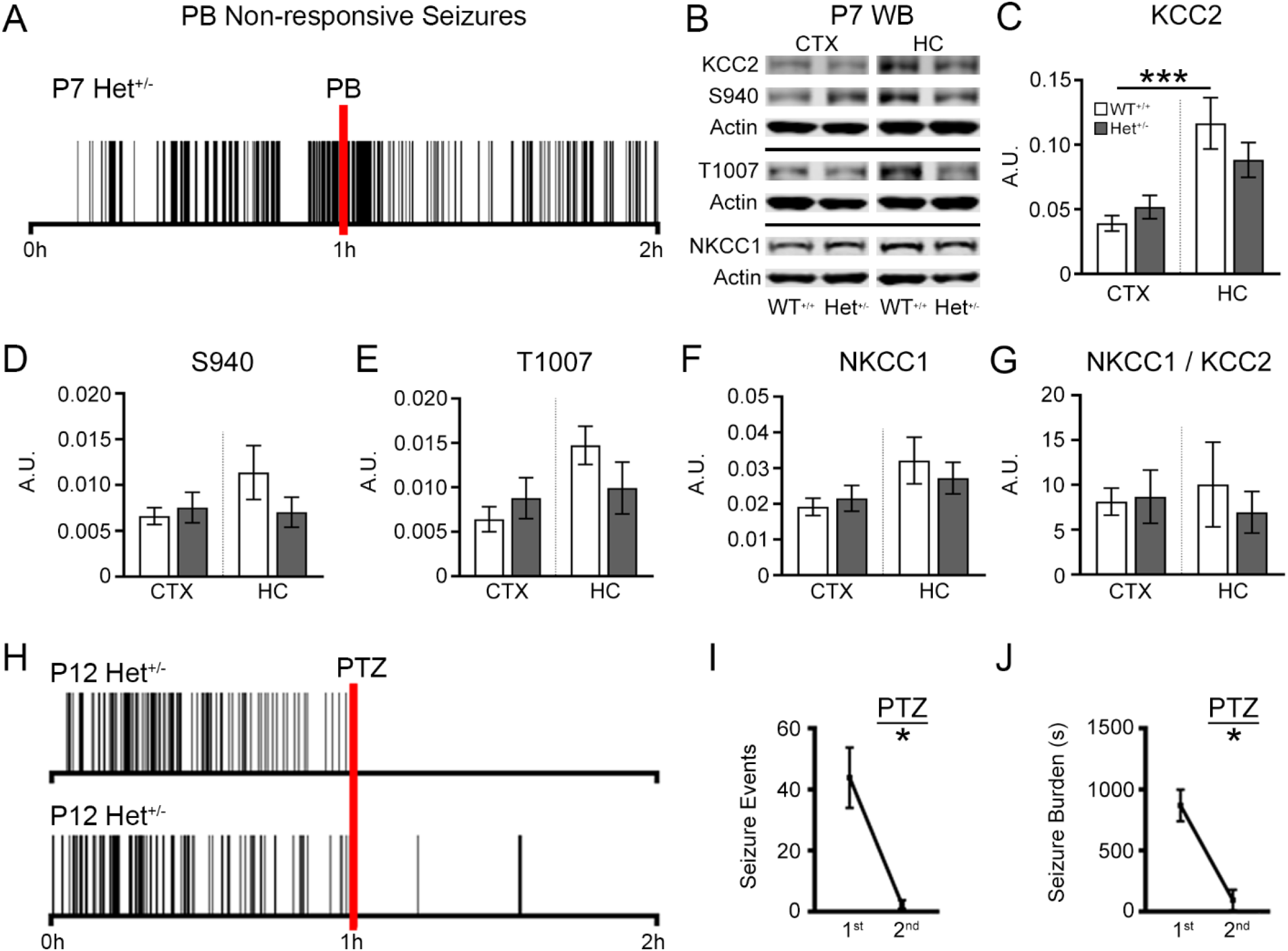
GABAergic signaling is associated with seizure activity in SYNGAP1^+/-^ mice. (**A**) Seizure frequency raster plot of a P7 Het mouse during 2h vEEG recording (n=2). Red bar represents a loading dose of PB (25mg/kg; IP injection). (**B)** Representative western blots showing KCC2, S940, T1007, and NKCC1 expression at P7 in WT and Het mice. All proteins of interest were normalized to β-actin. (**C**) KCC2 (n=12 samples per group), (**D**) S940 (n=6 samples per group), (**E**) T1007 (n=6 samples per group), and (**F**) NKCC1 expression in the cortex (CTX) and hippocampus (HC) of WT and Het mice (n=6 samples per group). (**G**) NKCC1 to KCC2 ratios for CTX and HC. WB results were gathered from n=3 mice per group. (**H**) Seizure frequency raster plot of P12 Het mice during 2h vEEG recording. Red bar represents a 20mg/kg dose of PTZ (IP injection). (**I**) 1^st^ and 2^nd^ h seizure events, and (**J**) 1^st^ and 2^nd^ h seizure burdens (n=3 P12 Het). (**C-G**) *P<0.05 and *P<0.001 by one-way ANOVA. (**I-J**) *P<0.05 by paired t-test.

### Gamma during NREM is high in juvenile Syngap1^+/-^ mice

In adult Het^+/-^ mice at P60 and P120, gamma power (35-50Hz) during NREM was higher than WT^+/+15^. It is currently unknown if the lack of behavioral state modulation in gamma power is present at younger ages. At P25, Het^+/-^ pups demonstrated high gamma power during NREM over 24h (Figure 4 A-B). Previously at P120, low dose PMP (2mg/kg intraperitoneal) reduced gamma power during NREM^15^. Low dose PMP restored cortical gamma behavioral-state modulation in juveniles Het^+/-^ pups (Figure 4 C), similar to previous results in adult Het^+/-^ mice^15^. To investigate if high gamma power during NREM is dependent upon sleep pressure, P35 mice underwent 6h of sleep deprivation during telemetric vEEG recording (Figure 4 D-E). Sleep deprivation significantly exacerbated high gamma power during NREM in P35 Hets^+/-^ (Figure 4 F-G). Importantly, no significant differences in NREM delta (0.5-4Hz) power were apparent between genotypes, a proxy for slow wave sleep compensation (Figure 4H).

**Figure 4.**
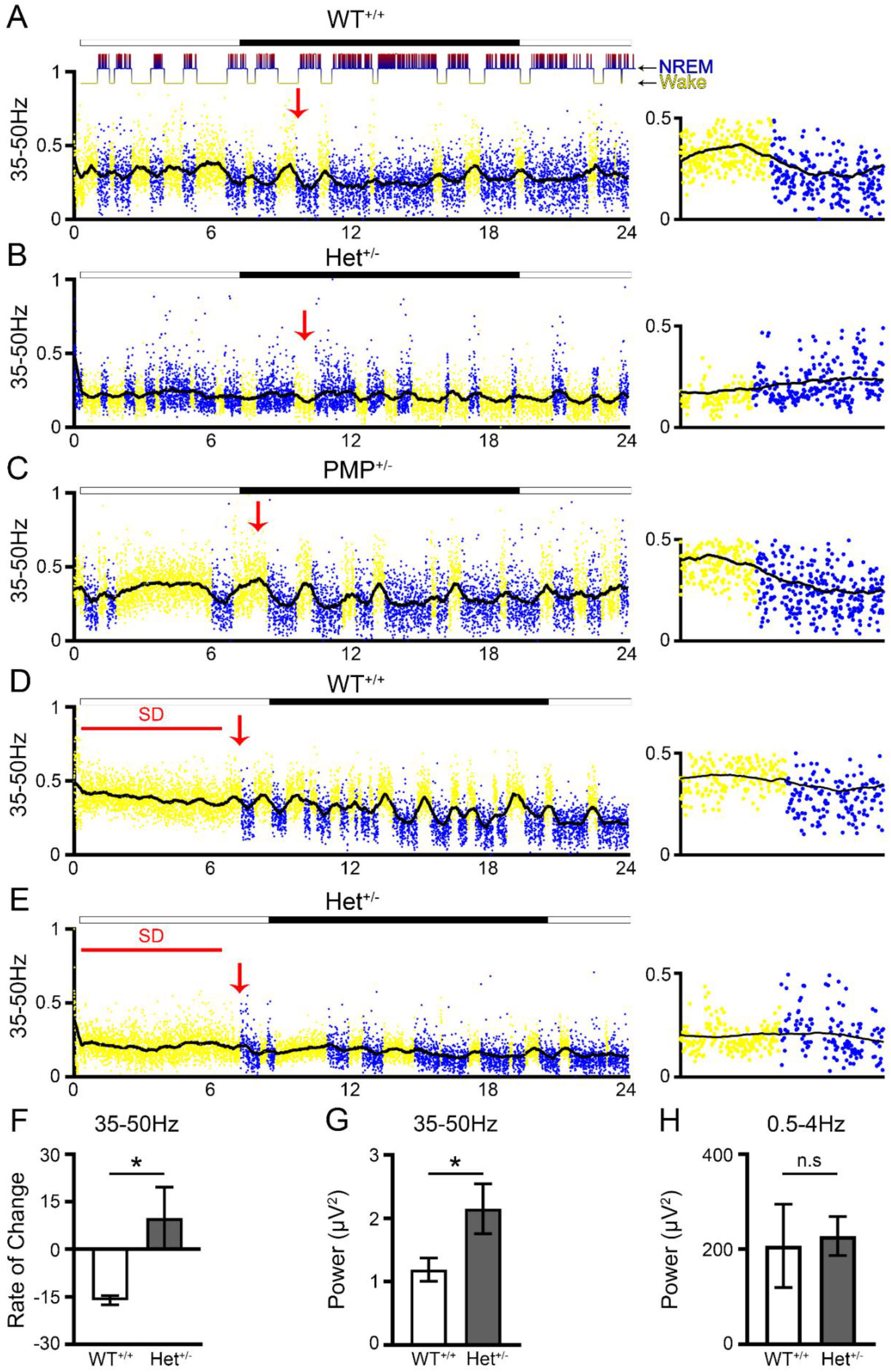
High Gamma Power during NREM in juvenile P24 Syngap1^+/-^ mice. (**A**) WT gamma (35-50Hz) trace for every 10 sec epoch over a 24h continuous EEG recording period demonstrates high gamma power during wake and low gamma power during NREM. Every dot represents gamma power for a 10s epoch. Yellow denotes wake-state, and blue denotes NREM sleep. The solid black line represents a running average. Hypnogram above graph A shows sleep and wake states for same mouse over 24h. Light cycles are depicted above graphs as lights on (white) or lights off (black). Red arrowheads show wake/sleep transition points for expanded timescale panels shown to the right. (**B**) Gamma power in juvenile P24 Het mice failed to transition to the lower NREM levels. (**C**) Low-dose PMP administration restored behavioral state-dependent gamma power in the same HET mice at P25. (**D**) 6h sleep deprivation (SD) during 24h-hour EEG recording in in WT and (**E**) Het mice. (**F**) Rate of change for gamma power during Wake to NREM transitions. (**G**) NREM Gamma power after 6h SD. (**H**) NREM Delta power (0.5-4Hz) after 6h SD.*P<0.05 two tailed t-test. (n=2 mice per group).

## Discussion

In this study, we found that *Syngap1* haploinsufficiency causes spontaneous recurrent seizures early in life (P7-P12). During development, persistent seizures and interictal epileptiform activity worsen developmental regression, cognitive function, and cause behavioral impairments^28,29^. Acute seizures alone, without a contributing genetic mutation, can disrupt thalamocortical critical period plasticity and are associated with impaired sensorimotor integration^30^. The phenotypic spectrum of *SYNGAP1* DEE may be strongly influenced by the interaction between the aberrant synaptic plasticity caused by *SYNGAP1* haploinsufficiency and the concurrent epilepsy^5,29^. Specifically, uncontrolled early-life seizures may compound the neurodevelopmental impairments caused by the lack of functional SYNGAP1. Our data suggests that early in life (P7), before SYNGAP1 reaches its peak in forebrain expression^10^, *Syngap1* haploinsufficiency results in aberrant epileptic circuits. Further, juvenile EEGs combined with sleep deprivation protocols identified the presence of impaired cortical gamma and the exacerbating effect of increased sleep pressure on this recently identified EEG biomarker.

### Early-life seizures in Het^+/-^ pups are electrographic and vary in frequency

Ongoing natural history studies in the growing *SYNGAP1* patient population are making it clear that epilepsy is commonly associated with *SYNGAP1* haploinsufficiency. Due to more awareness and early-life screening, seizures have been diagnosed in *SYNGAP1* patients as early as 4 months of life^5^. Our data indicates that many of these early-life seizures may require EEG for clinical diagnosis, as the majority of the seizures did not have an overt motor component. Subclinical seizures are a type of seizure that does not present any clinical signs or symptoms generally associated with seizures but show abnormal brain activity in the form of synchronous spike-wave discharges on EEG. Long term vEEG monitoring in developmental disorders associated with epilepsy can help capture these electrographic discharges. Our results suggest that early long-duration vEEG monitoring is warranted in all children with suspected pathogenic SYNGAP1 variants.

In our experiments, even standardized EEG recording durations on pups with identical mutations in *Syngap1* demonstrated wide variability in the incidence of early-life seizures in both sexes. This variability is clinically relevant as short vEEG monitoring may not be sufficient to identify the epilepsy. Our results allow for the future identification of susceptibility factors driving this variability in seizure frequency during development. As the genetic screenings in the future will inevitably start occurring earlier in postnatal life as part of genetic screening panels, vEEG recording will help classify the early-lie epilepsy in *SYNGAP1* DEE. An improved understanding of the epilepsy may help classify the phenotypic variability within *SYNGAP1* DEE patient cohorts. The contribution that early-life seizures have on the natural history of *SYNGAP1* DEE is an outstanding question of critical importance for future therapies. The further characterization of the early-life seizures in *Syngap1* DEE models will assist these endeavors.

### An emerging GABAergic hypothesis in *SYNGAP1* DEE

Our data suggests that GABAergic signaling during development promotes ictogenesis in *Syngap1* DEE. A loading dose of PB, a positive allosteric modulator of GABA_A_Rs, failed to curb P7 seizures. In contrast, the GABA_A_R antagonist PTZ significantly reduced seizures at P12. These findings suggest that GABAergic signaling may contribute to early-life seizure generation in *Syngap1* DEE. In mature neurons both the neuronal Cl^−^ gradient and efficacy of GABA_A_-mediated synaptic transmission is influenced by the Cl^−^ exporter KCC2 maintaining a low intracellular Cl^−^ concentration ([Cl^−^]_i_)^25^. Early in brain development, KCC2 expression is low and [Cl^−^]_i_. is high, resulting in depolarizing GABAergic signaling^31^. In excitotoxic conditions, KCC2 hypofunction may facilitate the emergence of refractory seizures^23,32^. ~50% of *SYNGAP1* children are known to have refractory seizures^533^. Here, we report refractoriness to a first-line positive GABA_A_R modulator at P7. However, KCC2 expression and phosphorylation levels were not found to be deficient nor was the NKCC1 expression high. These findings indicate that Cl^−^ cotransporter functional deficits may not play a role in the unique drug responses reported here. Previous studies have identified increased synaptic inhibition during development in Het^+/-^ mice^13,20^, supporting the results reported here.

*Syngap1* has been shown to play a critical role in GABAergic circuit development and function^16,34^. Previously in adult Het^+/-^ mice our group identified an increased expression in the calcium impermeable AMPA subunit GluA2 in PV interneurons and disrupted behavioral-dependent gamma oscillations^15^. Here we document the early emergence of abnormal behavioral-dependent gamma oscillations with no evidence of spontaneous seizures at P24-P40. This may indicate that abnormal gamma is independent of seizure occurrence and represents an underlying circuit dysfunction. Future studies will delineate the impact of early life seizures on these novel biomarkers.

### Increasing sleep pressure further aggravates abnormal cortical gamma

During wake, experience dependent plasticity strengthens excitatory neuronal synapses that permit the storage of information. Sleep is a necessary behavior that allows neurons to consolidate information and permits synaptic renormalization^35,36^. Sleep-wake patterns and alertness level during wakefulness are known to be modulated by two interacting processes: one is the sleep pressure that increases as a saturating exponential during wakefulness; the second are the circadian circuits in the brain that drive the internal oscillatory rhythm that run our 24h cycles^36^. Numerous studies have documented slow wave sleep compensation and its characteristics following sleep deprivation. Additionally, the detrimental effects of prolonged sleep deprivation on cognitive performance are also well established^37–40^. Our previous study established significant disruption of cortical gamma homeostasis in adult Het^+/-^ mice^15^. The identification of normal slow wave sleep compensation to the increased sleep pressure in both WT^+/+^ and Het^+/-^ pups highlight the uniqueness of the qEEG biomarker related to gamma homeostasis which is known to depend on function of fast-spiking PV interneurons^41,42^. Significant aggravation of cortical gamma homeostasis during NREM following increase in sleep pressure further uncovers the role of PV dysfunction in circuits governing sleep homeostasis.

## Conclusion

For the first time in a *Syngap1* encephalopathy mouse model, we have identified the occurrence of early-life seizures. The anti-seizure response to a GABA_A_R antagonist suggests a critical role for GABAergic signaling in early ictogenesis. Any prospective evidence-based therapies will have to consider the effect of repeated seizures during development on the pathophysiology of the *SYNGAP1* DEE.

## Supporting information

Video suppl for Fig, 2D

## Acknowledgement

Research reported in the publication was supported by the Eunice Kennedy Shriver National Institute of Child Health and Human Development of National Institutes of Health under Grant No. R01HD090884 (SDK). The content is solely the responsibility of the authors and does not necessarily represent the official view of the NIH. We thank Dr. Richard Huganir’s Lab for generously helping us establish our own *Syngap1* mouse colony, specifically Dr. Ingie Hong and Dr. Yoichi Araki.

## Author Contributions

SDK conceived the project. BJS, PAK, and SDK acquired data. BJS, PAK, SGA, and SDK analyzed data. BJS, PAK, and SDK wrote the paper.

## Abbreviations

(SynGAP1): Synaptic Ras GTPase-activating protein 1
(PB): Phenobarbital
(PTZ): Pentylenetetrazol
(KCC2): K-Cl cotransporter 2
(NKCC1): Na-K-Cl cotransporter 1
(GABA_A_R): GABA_A_ receptor
(ASMs): Anti-seizure Medications

---

**Supplemental Figure 1.**
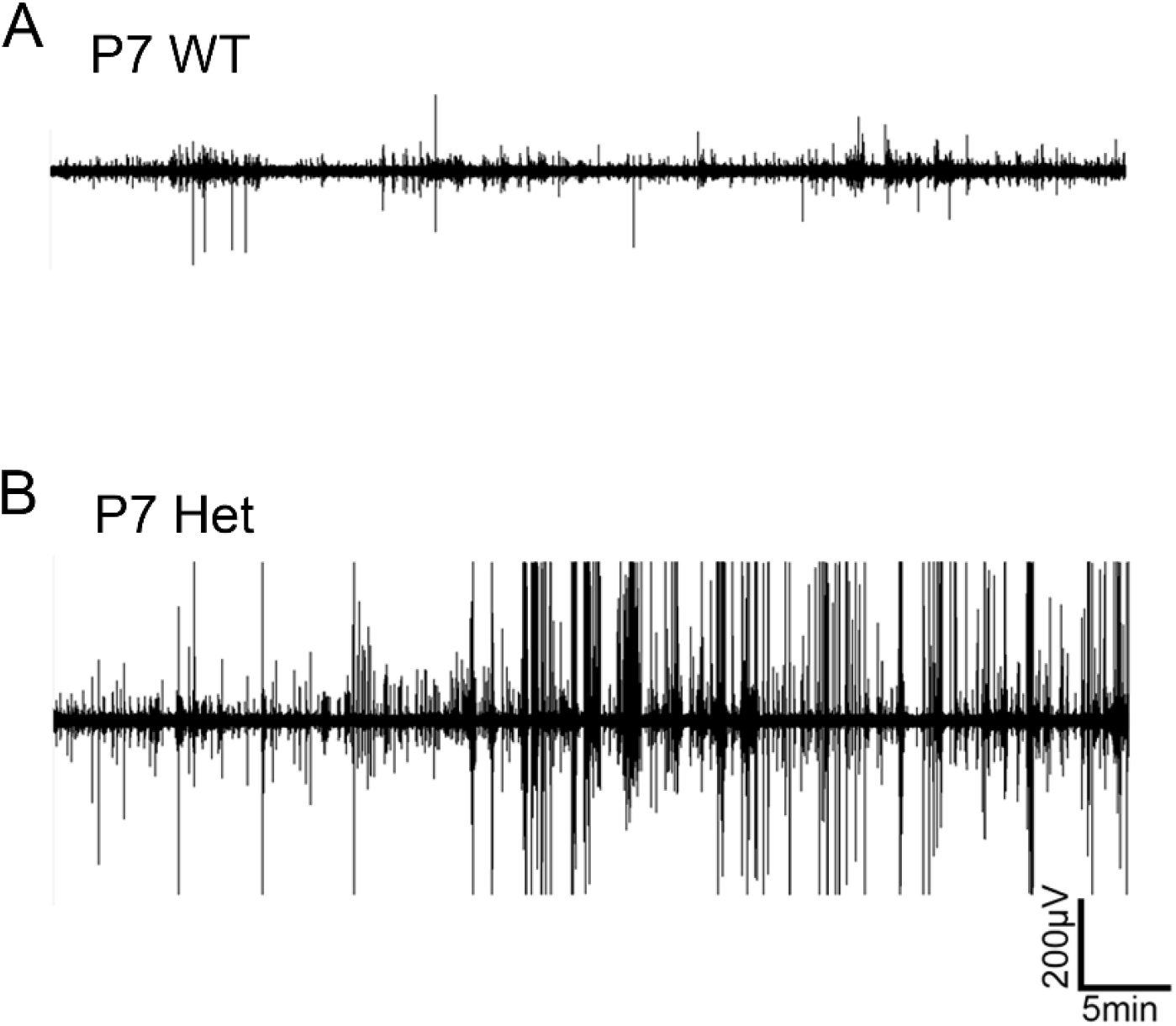
1h EEG trace showing burden of spontaneous seizures in SYNGAP1^+/-^ mice. (**A**) 1h WT and (**B)** Het trace at P7.

**Supplemental Table 1.**

| Reagent type (species) or resource | Designation | Source Reference | Identifiers | Additional Information |
| --- | --- | --- | --- | --- |
| Genetic reagent (M. musculus) | B6.129-Syngap1<sup>tm1Rlh/J</sup> | Jax | RRID:IMSR_JAX:008890 | Dr. Richard L Haganir, Johns Hopkins University |
| Chemical compound, drug | Phenobarbital (PB) | MilliporeSigma | P5178 | N/A |
| Chemical compound, drug | Pentylene-tetrazol (PTZ) | MilliporeSigma | P6500 | N/A |
| Chemical compound, drug | DMSO | Sigma | D8418 | N/A |
| Software, algorithm | Graphpad Prism | Graphpad Software | RRID:SCR_002798 | 8 |
| Software, algorithm | Sirenia | Pinnacle Technology | pinnaclet.com/sirenia | 3-Channel EEG/EMG Tethered Mouse System |
| Antibody | mouse $\alpha$ KCC2 | Aviva Systems Biology OASE00240 | AB_2721238 | 1:1000; WB |
| Antibody | rabbit $\alpha$ pKCC2-S940 | Aviva Systems Biology OAPC00188 | AB_2721198 | 1:1000; WB |
| Antibody | rabbit $\alpha$ pKCC2-1007 | PhosphoSolutions p1551-1007 | AB_2716769 | 1:1000; WB |
| Antibody | rabbit $\alpha$ NKCC1 | Sigma-Aldrich AB3560P | AB_91514 | 1:1000; WB |
| Antibody | mouse $\alpha$ actin | LI-COR Biosciences 926-42213 | AB_2637092 | 1:10000; WB |
| Antibody | goat $\alpha$ mouse IgG, IRDye® 800CW Conjugated | LI-COR Biosciences 926-32210 | AB_621842 | 1:5000; WB |
| Antibody | goat $\alpha$ rabbit IgG Antibody, IRDye® 680LT Conjugated | LI-COR Biosciences 926-68021 | AB_10706309 | 1:5000; WB |

